# Raybloc^®^: A Marine Bioactive Silica-Microsponge Formulation Confers Superior Protection against Blue Light and Infrared-A Induced Skin Damage in Murine Model

**DOI:** 10.64898/2026.03.21.713389

**Authors:** Shaylynn Yu, Kevin Ngo, Muhammad Ovais

## Abstract

Long-term exposure to high-energy visible (HEV) blue light and infrared-A (IR-A) radiation accelerates oxidative stress, inflammation, and transepidermal water loss (TEWL), leading to photoaging and damage to the skin barrier. In this study, we developed Raybloc®, a marine bioactive silica microsponge formulation, and evaluated its protective effects against combined high-energy visible (HEV; 410–480 nm) and infrared-A (IR-A; 700–1400 nm) exposure in a preclinical model. We divided 36 nude BALB/c-nu/nu mice into six groups: one that didn’t get any treatment, one that got Raybloc® (no radiation), one that got Raybloc® 5%, one that got Raybloc® 8%, one that got HA 0.5%, and one that got HA 0.8%. Animals underwent topical treatment for 14 days under regulated exposure to HEV (410–480 nm, 100 J/cm^2^/day) and IR-A (700–1400 nm, 30 mW/cm^2^). We examined transepidermal water loss (TEWL), skin hydration, oxidative stress, inflammatory cytokines (IL-1β, IL-6, TNF-α, IL-10), and histological indicators of collagen preservation through biophysical, biochemical, and histopathological techniques. In the Raybloc® 8% group, TEWL dropped by 48.3 ± 4.6% (p < 0.001), and skin hydration went up by 62.7 ± 5.1%. The levels of ROS and MMP-1 expression decreased by 63.4% and 57.2%, respectively, while collagen I increased by 2.1 times compared to HA 0.8%. There was a big drop in the pro-inflammatory cytokines IL-1β, IL-6, and TNF-α (−54%, −49%, and −46%), and a big rise in IL-10 (+38%). Histological analysis demonstrated well-preserved epidermal integrity and dense collagen bundles in Raybloc®-treated mice, whereas irradiated controls exhibited dermal disorganization and inflammatory infiltration. Raybloc® showed better photoprotective, antioxidant, and moisturizing effects than HA-based products. It also helped reduce oxidative and inflammatory skin damage caused by blue light and IR-A. These results support Raybloc® as a next-generation multifunctional dermocosmetic that can help stop photoaging caused by digital and solar radiation.

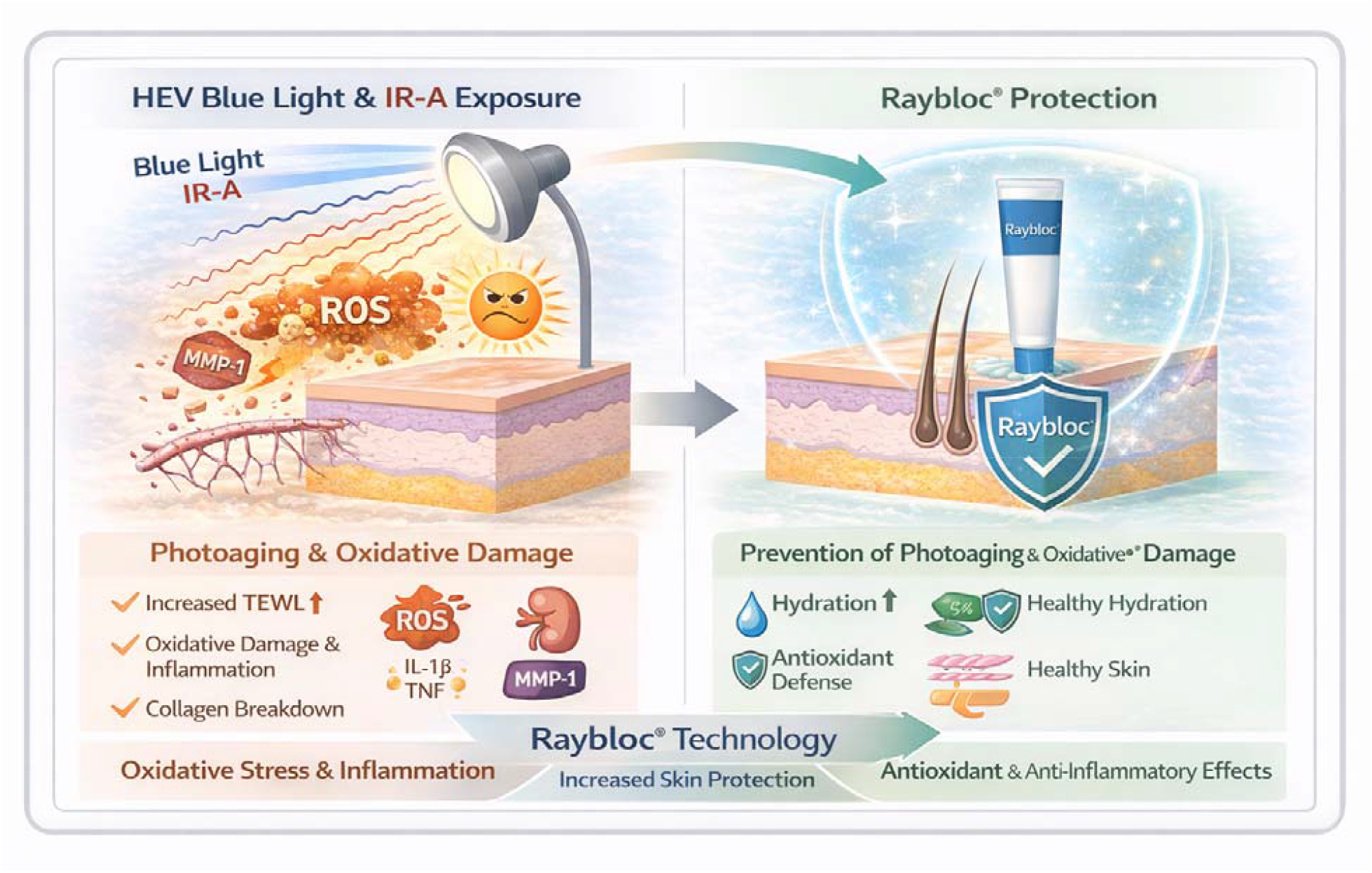

## 1. Introduction

The skin is the body’s main barrier and is exposed to different environmental stressors, such as high-energy visible (HEV) blue light (400–490 nm) and infrared-A (IR-A) radiation (700–1400 nm). Ultraviolet (UV) radiation has long been acknowledged for its photoaging effects; however, increasing evidence suggests that high-energy visible (HEV) and infrared-A (IR-A) radiation also elicit similar oxidative and inflammatory responses that impair skin structure and function (Gromkowska-Kępka, Puścion - Jakubik, Markiewicz-Żukowska, & Socha, 2021; Salminen, Kaarniranta, & Kauppinen, 2022). Long-term exposure leads to higher levels of reactive oxygen species (ROS), mitochondrial damage, collagen breakdown, and a weakened epidermal barrier, which can manifest as dry skin, reduced elasticity, and accelerated aging (Dupont, Gomez, & Bilodeau, 2013; Lan, 2019).

On the other hand, HEV and IR-A radiation penetrate deeper into the dermis and alter fibroblast function and extracellular matrix assembly (Garimano, Aguayo Frías, & González Maglio, 2025). These wavelengths trigger the release of pro-inflammatory cytokines (IL-1β, IL-6, TNF-α), activate the NF-κB and MAPK pathways, and increase the levels of matrix metalloproteinases (MMPs), such as MMP-1, which accelerate collagen breakdown (Pinto et al., 2020). To protect against HEV and IR-A, we need antioxidants, anti-inflammatories, and moisturizers that can lower ROS levels and keep the skin hydrated. Marine-derived bioactives, such as extracts from *Chlorella vulgaris* and *Laminaria japonica*, comprise peptides, polysaccharides, and polyphenols that exhibit significant antioxidant and anti-photoaging properties (Prates, 2025). Peptides from chlorella can boost the body’s own enzymes that protect it (superoxide dismutase, catalase, and glutathione peroxidase). Polysaccharides from Laminaria can also reduce MMP activity and help the body produce more collagen (Hu et al., 2016). Glycerin and sorbitol are two examples of humectants that help the stratum corneum retain water and reduce transepidermal water loss (TEWL) (Lodén, 2003).

Microsponge technology enables controlled release of encapsulated components, thereby enhancing their photostability and bioavailability (Abdalla, Osman, Nouh, & El Maghraby, 2021; Jadhav et al., 2013; Rahman et al., 2022). This study examined the protective and moisturizing efficacy of Raybloc® (marine bioactive silica-microsponge formulation) against blue light- and IR-A-induced skin damage in a nude mouse model, using hyaluronic acid (HA) as a hydration comparator. We proposed that Raybloc®, through its synergistic combination of marine antioxidants, humectants, and microsponge delivery, would markedly reduce oxidative stress, inflammation, and transepidermal water loss (TEWL), whilst enhancing collagen integrity and hydration. Unlike sunscreens, which are designed to attenuate ultraviolet radiation through optical filtering and SPF-based mechanisms, dermocosmetic photoprotective ingredients aim to biologically mitigate light-induced oxidative stress, inflammation, and barrier dysfunction.

## 2. Materials and Methods

### 2.1. Materials

Raybloc® is a silica-derived microcarrier formulation platform loaded with marine-derived bioactive agents (Chlorella extract and Laminaria extract, along with humectants such as glycerin and sorbitol) dispersed within a stabilized aqueous microfluid system (proprietary information provided under NDA). Hyaluronic acid (HA) gels at 0.5% and 0.8% concentrations were used as comparators. Reagents for biochemical assays, including ELISA kits for IL-1β, IL-6, TNF-α, IL-10, and ROS and MMP-1 quantification, were from Thermo Fisher Scientific. We got H&E, Masson’s Trichrome, and immunohistochemistry reagents from Sigma-Aldrich.

### 2.2. Animal Model and Experimental Design

Thirty-six female BALB/c-nu/nu nude mice (6–8 weeks old, 20–25 g) were sourced from the National Institute of Health, Islamabad, Pakistan, and maintained in standard conditions (22 ± 2 °C, 50 ± 5% humidity, 12-h light/dark cycle) with unrestricted access to food and water. The Institutional Animal Ethics Committee approved all procedures with reference No. 9812, which followed the OECD and ARRIVE 2.0 guidelines. Mice were randomly allocated into six groups (n = 6/group):

| GROUP | TREATMENT | LIGHT EXPOSURE | PURPOSE |
| --- | --- | --- | --- |
| G1 | Untreated | None | Negative control (baseline) |
| G2 | Raybloc® (8%) | None | Moisturization/safety control |
| G3 | Raybloc® (5%) | Blue + IR-A | Test group (low-dose) |
| G4 | Raybloc® (8%) | Blue + IR-A | Test group (high-dose) |
| G5 | HA (0.5%) | Blue + IR-A | Comparator (low-dose HA) |
| G6 | HA (0.8%) | Blue + IR-A | Comparator (high-dose HA) |

Treatments (100 µL) were applied topically to a 3 × 3 cm shaved area of the dorsal skin once daily for 14 days.

### 2.3. Light Exposure Protocol

A calibrated LED array sent blue light (410–480 nm, 100 J/cm^2^/day), and a filtered halogen lamp made IR-A radiation (700–1400 nm, 30 mW/cm^2^). The distance between the light source and the skin was kept at 15 cm. During days 4–10, mice were exposed to blue light for 30 minutes and IR-A for 20 minutes every day. They were treated for three days before treatment and three days after treatment.

### 2.4. Measurement of Skin Barrier Function

#### 2.4.1. Transepidermal Water Loss (TEWL)

We used a Tewameter® TM300 (Courage + Khazaka, Germany) to measure transepidermal water loss (TEWL), a marker of the skin barrier’s health. Before each measurement session, the animals were acclimatized to the testing environment for 20-30 minutes at a controlled temperature (22 ± 2 °C) and humidity (45–55%) to minimize environmental effects. The probe was carefully placed at a right angle to the skin and held still for 10 to 15 seconds before recording. Three readings were taken from each site, one after the other, and the average was used to reduce differences between individuals. Measurements were taken on days 0, 3, 7, 10, and 14 at the specified dorsal application site.

#### 2.4.2. Skin Hydration

We used a Corneometer® CM825 (Courage + Khazaka, Germany) to determine the moisture content of the stratum corneum. This device works by measuring changes in capacitance. Before the assessment, the skin surface was meticulously cleansed with sterile saline and allowed to equilibrate for 5 minutes to prevent artifacts caused by residual moisture. At each time point, three measurements were taken at the same anatomical location. The probe pressure and angle stayed the same for all of them. On days 0, 3, 7, 10, and 14, we checked and reported hydration levels in arbitrary hydration units (a.u.). Higher numbers meant that the skin could hold more water.

### 2.5. Histological and Immunohistochemical Analysis

On the 14th day, full-thickness skin samples were collected from both the treated and control areas. They were then placed in 10% neutral-buffered formalin for 24 to 48 hours to preserve tissue structure. After being fixed, samples were dried out with a series of graded ethanol, then cleared with xylene, and finally put in paraffin. Using a rotary microtome, paraffin blocks were sectioned at 5 µm. These sections were then mounted on glass slides coated with poly-L-lysine to ensure they stuck well. For general histomorphological assessment, sections were stained with hematoxylin and eosin (H&E). Epidermal and dermal structures were scrutinized for alterations in epidermal thickness, inflammatory cell infiltration, stratum corneum integrity, and dermal tissue remodeling.

To prepare paraffin sections for immunohistochemical analysis (IHC), they were first soaked in xylene and then in decreasing concentrations of ethanol. We used a water bath at 95–98 °C for 20 minutes to recover the antigen from citrate buffer (pH 6.0). We used 3% hydrogen peroxide to stop endogenous peroxidase activity and 5% bovine serum albumin (BSA) to stop nonspecific binding. Then, the sections were incubated overnight at 4 °C with primary antibodies against IL-1β and TNF-α at a 1:200 dilution. The next day, slides were incubated with a secondary antibody conjugated to HRP and developed with DAB (3,3′-diaminobenzidine) and stained with hematoxylin.

### 2.6. Oxidative Stress and Molecular Biomarkers

We examined oxidative stress markers and molecular signaling pathways using biochemical techniques, quantitative PCR, and immunoassays. On the 14th day, skin tissues were quickly frozen in liquid nitrogen and stored at −80 °C until they were ready for inspection. For homogenate preparation, samples were weighed and mixed in ice-cold PBS containing protease and phosphatase inhibitor cocktails. Homogenates were centrifuged at 12,000 × g for 15 minutes at 4 °C, and the supernatants were used for subsequent biochemical tests.

We used the 2’,7’-dichlorofluorescin diacetate (DCFH-DA) fluorescence method to determine the amount of reactive oxygen species (ROS) present. In brief, we incubated small amounts of tissue lysate in the dark with 10 µM DCFH-DA at 37 °C for 30 minutes. The non-fluorescent DCFH-DA is converted to the fluorescent DCF when ROS are present in the cell. A microplate reader measured fluorescence intensity at 485/530 nm.

For quantitative real-time PCR (qPCR), total RNA was isolated from skin tissues using TRIzol® reagent according to the manufacturer’s protocol. We used a NanoDrop spectrophotometer to see how pure and concentrated the RNA was. We used a high-capacity cDNA synthesis kit to generate complementary DNA (cDNA). We did qPCR with a real-time PCR system and SYBR Green master mix. We developed primers that worked with the genes Nrf2, HO-1, MMP-1, and Collagen I. GAPDH was the internal housekeeping gene that maintained steady expression levels. The amplification conditions included an initial denaturation step, followed by 40 cycles of denaturation, annealing, and extension. Gene expression levels of Nrf2, HO-1, MMP-1, and Collagen I were quantified by qPCR and expressed as fold change using the 2^^-ΔΔCt^ method. We also used ELISA kits to measure cytokine levels (IL-1β, IL-6, TNF-α, and IL-10) according to the instructions provided with the kits. At 450 nm, the absorbance was measured, and standard curves were used to fill in the gaps and find the concentrations.

### 2.7. Statistical Analysis

All experimental data were presented as mean ± standard deviation (SD). One-way analysis of variance (ANOVA) was employed to compare groups when the conditions of normality and homogeneity of variance were met. We used Tukey’s multiple-comparison post hoc test to find differences between groups after getting significant ANOVA results.

## 3. Results

### 3.1. Effect of Raybloc® on Skin Barrier Function

#### 3.1.1. Transepidermal Water Loss (TEWL)

Raybloc® treatment significantly decreased TEWL compared with irradiated controls and HA comparators during the study period **(Table 1)**. On day 14, TEWL values were 11.3 ± 0.9 g/m^2^/h in the Raybloc® 8% group and 21.8 ± 1.2 g/m^2^/h in the untreated irradiated control (p < 0.001). The 5% formulation resulted in a 35% decrease, whereas the 0.8% HA formulation yielded merely a 21% decrease.

**Table 1.**
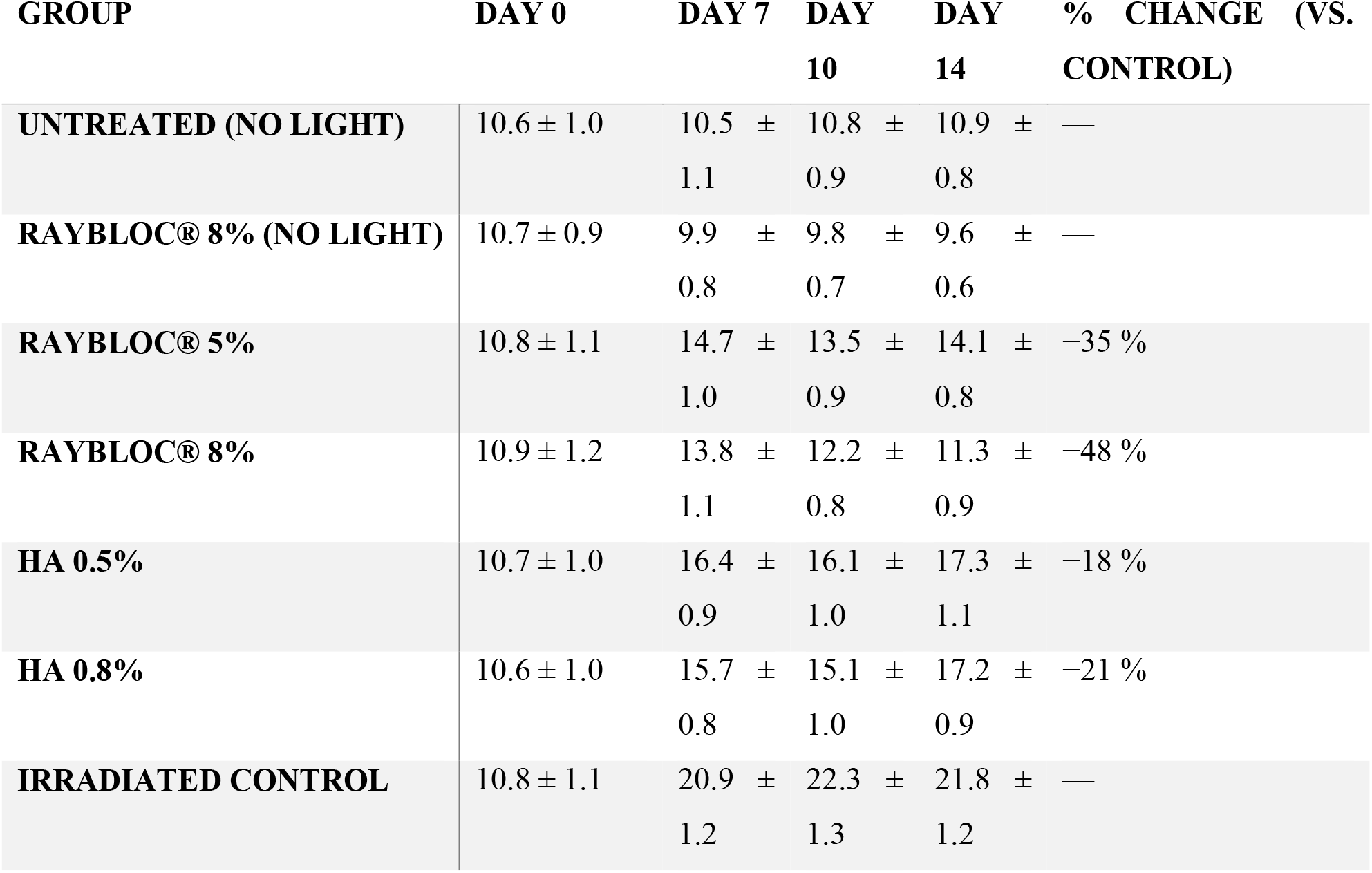
Transepidermal water loss (TEWL) measurements across study groups.

| GROUP | DAY 0 | DAY 7 | DAY 10 | DAY 14 | % CHANGE (VS. CONTROL) |
| --- | --- | --- | --- | --- | --- |
| UNTREATED (NO LIGHT) | $10.6 \pm 1.0$ | $10.5 \pm 1.1$ | $10.8 \pm 0.9$ | $10.9 \pm 0.8$ | — |
| RAYBLOC® 8% (NO LIGHT) | $10.7 \pm 0.9$ | $9.9 \pm 0.8$ | $9.8 \pm 0.7$ | $9.6 \pm 0.6$ | — |
| RAYBLOC® 5% | $10.8 \pm 1.1$ | $14.7 \pm 1.0$ | $13.5 \pm 0.9$ | $14.1 \pm 0.8$ | –35 % |
| RAYBLOC® 8% | $10.9 \pm 1.2$ | $13.8 \pm 1.1$ | $12.2 \pm 0.8$ | $11.3 \pm 0.9$ | –48 % |
| HA 0.5% | $10.7 \pm 1.0$ | $16.4 \pm 0.9$ | $16.1 \pm 1.0$ | $17.3 \pm 1.1$ | –18 % |
| HA 0.8% | $10.6 \pm 1.0$ | $15.7 \pm 0.8$ | $15.1 \pm 1.0$ | $17.2 \pm 0.9$ | –21 % |
| IRRADIATED CONTROL | $10.8 \pm 1.1$ | $20.9 \pm 1.2$ | $22.3 \pm 1.3$ | $21.8 \pm 1.2$ | — |

#### 3.1.2. Skin Hydration

Corneometer® measurements of hydration levels showed a significant improvement in the Raybloc®-treated groups **(Table 2)**. After 14 days, hydration increased by 62.7 ± 5.1% in the Raybloc® 8% group compared to irradiated controls (p < 0.001), exceeding both HA comparators.

**Table 2.** Corneometer® hydration readings (arbitrary units).

| GROUP | DAY 0 | DAY 7 | DAY 10 | DAY 14 | % INCREASE (VS. CONTROL) |
| --- | --- | --- | --- | --- | --- |
| UNTREATED (NO LIGHT) | 38.1 ± 2.0 | 39.2 ± 2.2 | 40.1 ± 2.1 | 41.3 ± 2.0 | — |
| RAYBLOC® 8% (NO LIGHT) | 38.5 ± 1.9 | 44.8 ± 2.3 | 47.5 ± 2.4 | 49.2 ± 2.5 | — |
| RAYBLOC® 5% | 38.0 ± 2.0 | 42.9 ± 2.1 | 45.1 ± 2.2 | 46.7 ± 2.3 | +48 % |
| RAYBLOC® 8% | 38.4 ± 2.1 | 44.2 ± 2.3 | 47.8 ± 2.4 | 49.9 ± 2.6 | +63 % |
| HA 0.5% | 38.2 ± 2.0 | 41.0 ± 2.1 | 42.3 ± 2.2 | 43.6 ± 2.1 | +28 % |
| HA 0.8% | 38.1 ± 2.0 | 42.7 ± 2.1 | 44.3 ± 2.3 | 45.2 ± 2.4 | +36 % |
| IRRADIATED CONTROL | 38.3 ± 2.1 | 31.4 ± 2.0 | 29.5 ± 1.8 | 30.7 ± 1.9 | — |

### 3.2. Histopathological and Immunohistochemical Evaluation

H&E staining showed that irradiated controls exhibited marked epidermal hyperplasia, spongiosis, and inflammatory infiltration **(Figure 1)**. In contrast, the Raybloc® groups had almost normal epidermal morphology and less immune-cell infiltration. Masson’s trichrome staining showed that Raybloc® 8% (collagen density = 85 ± 4%) retained more collagen than irradiated controls (42 ± 6%, p < 0.001). HA 0.8% achieved moderate collagen preservation (64 ± 5%).

**Figure 1.**
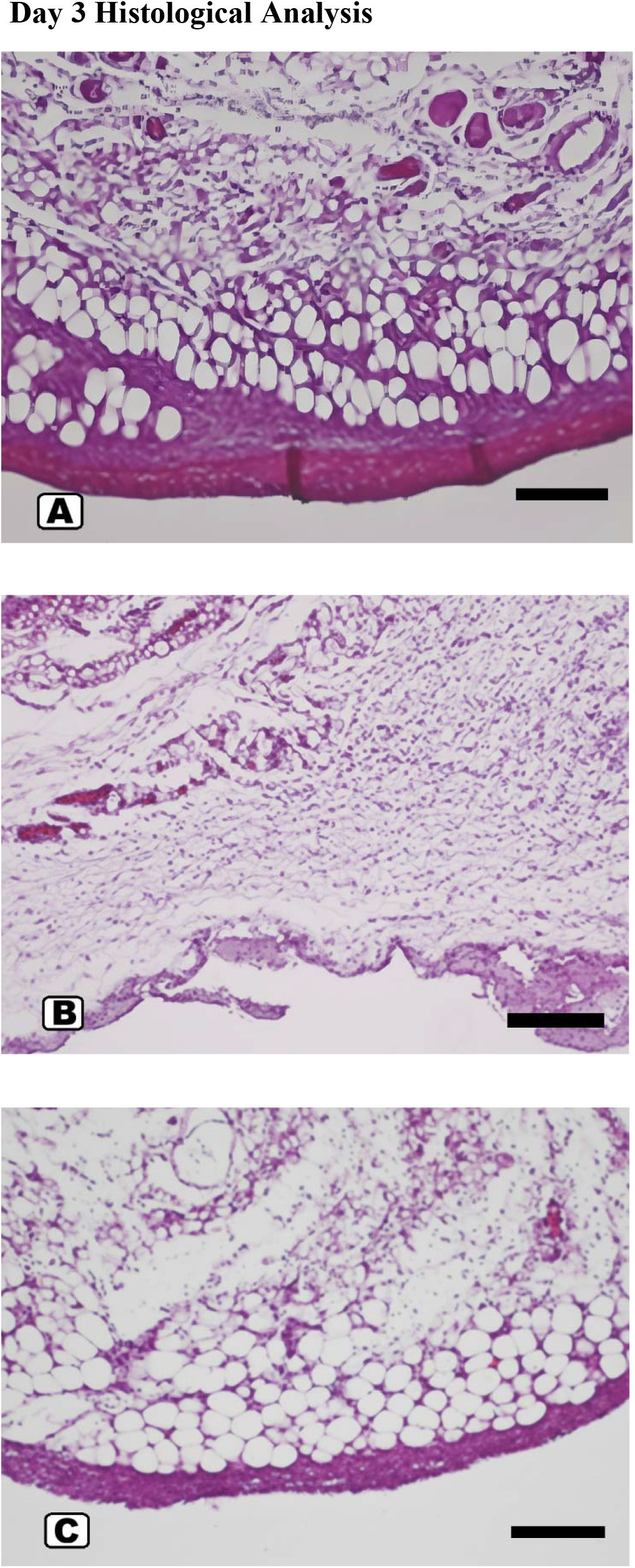
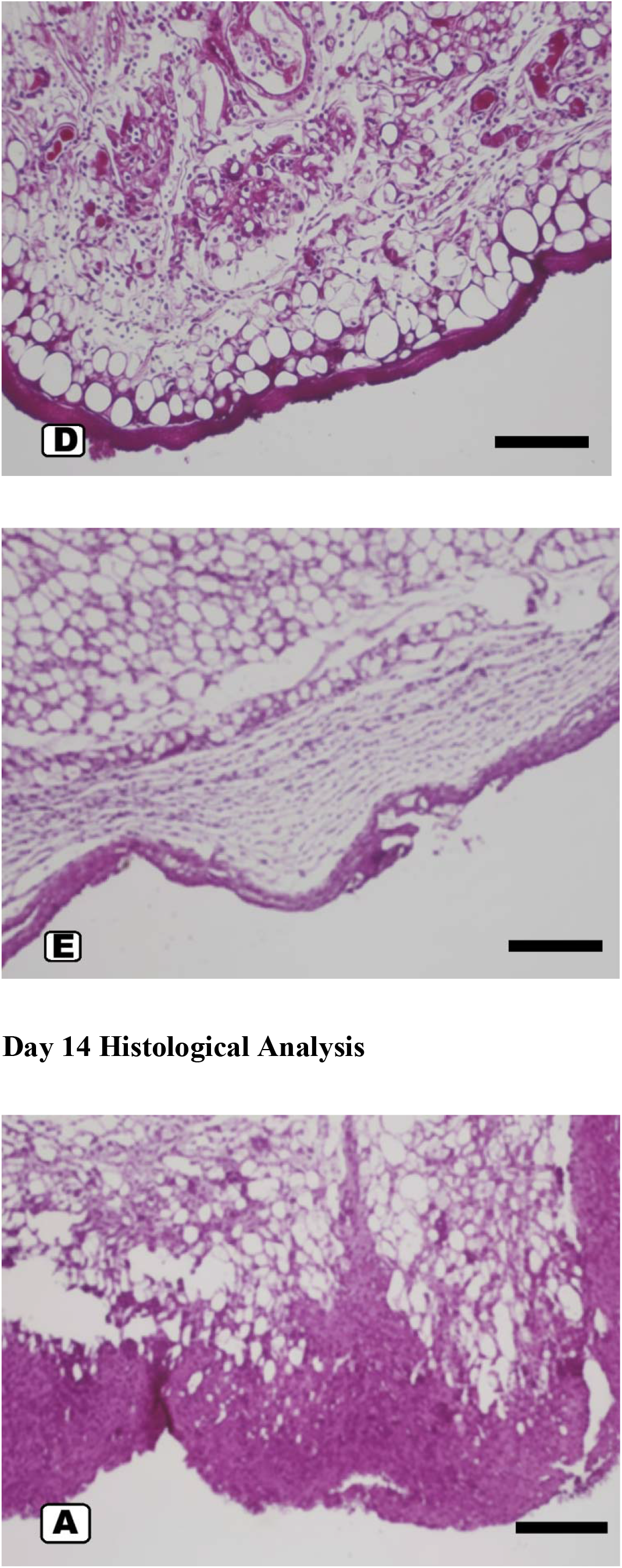
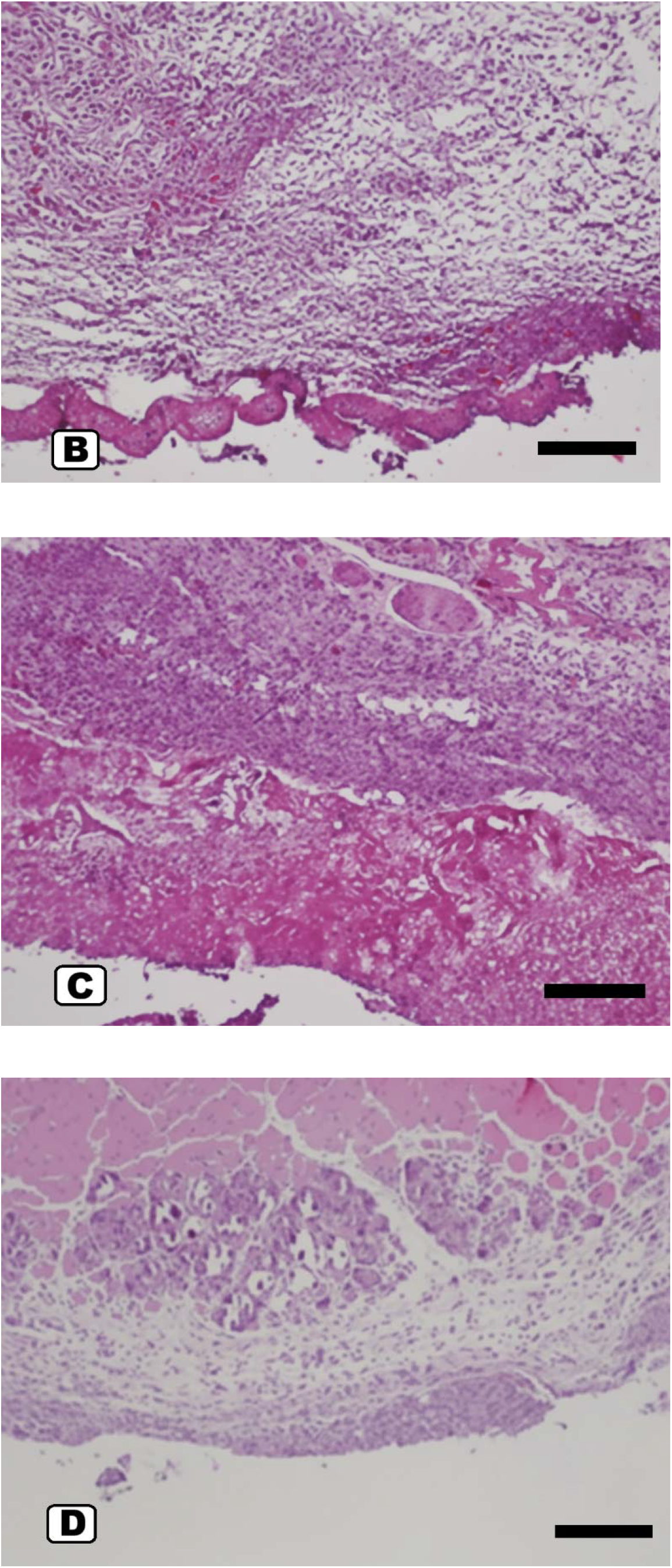
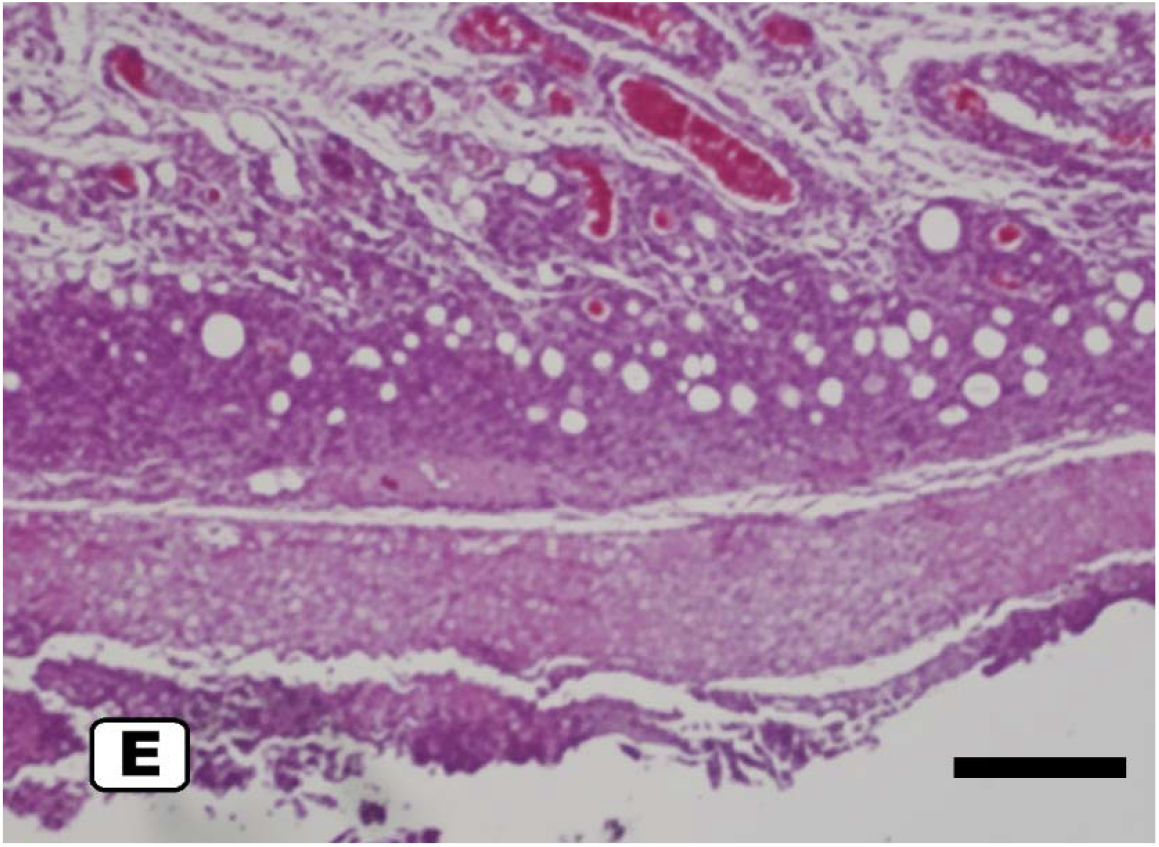
Histological comparison of epidermal thickness and collagen organization on Day 3 and Day 14. Raybloc® treatment preserved dermal collagen and reduced epidermal thickening. Scale bar, 100 μm.

### 3.3. Molecular and Biochemical Markers

Raybloc® reduced oxidative and inflammatory stress caused by HEV + IR-A exposure **(Table 3)**. The levels of ROS and MMP-1 were significantly lower, while those of Nrf2 and HO-1 were higher, indicating a stronger antioxidant response.

**Table 3.** Oxidative stress and gene expression analysis of antioxidant and photoaging markers (fold change relative to untreated control).

| MARKER | IRRADIATED<br>CONTROL | HA<br>0.8% | RAYBLOC®<br>5% | RAYBLOC®<br>8% |
| --- | --- | --- | --- | --- |
| ROS (RELATIVE<br>FLUORESCENCE, EXPRESSED<br>AS FOLD CHANGE) | 3.6 ± 0.3 × | 2.8 ± 0.2 × | 1.8 ± 0.2 × | 1.3 ± 0.1 × |
| NRF2 (MRNA FOLD) | 0.6 ± 0.1 | 1.2 ± 0.1 | 1.7 ± 0.1 | 2.3 ± 0.2 |
| HO-1 (MRNA FOLD) | 0.7 ± 0.1 | 1.4 ± 0.2 | 2.0 ± 0.1 | 2.6 ± 0.2 |
| **MMP-1 (MRNA FOLD)** | 4.1 ± 0.4 | 2.7 ± 0.3 | 1.9 ± 0.2 | 1.3 ± 0.2 |
| **COLLAGEN I (MRNA FOLD)** | 0.4 ± 0.1 | 1.2 ± 0.1 | 1.7 ± 0.2 | 2.1 ± 0.2 |

### 3.4. Inflammatory Cytokine Profile

ELISA quantification showed that pro-inflammatory cytokines were greatly reduced in the Raybloc® groups **(Table 4)**. Raybloc® 8% lowered the levels of IL-1β, IL-6, and TNF-α by 54%, 49%, and 46%, respectively, and raised the level of IL-10 by 38% compared to controls that had been irradiated.

**Table 4.**
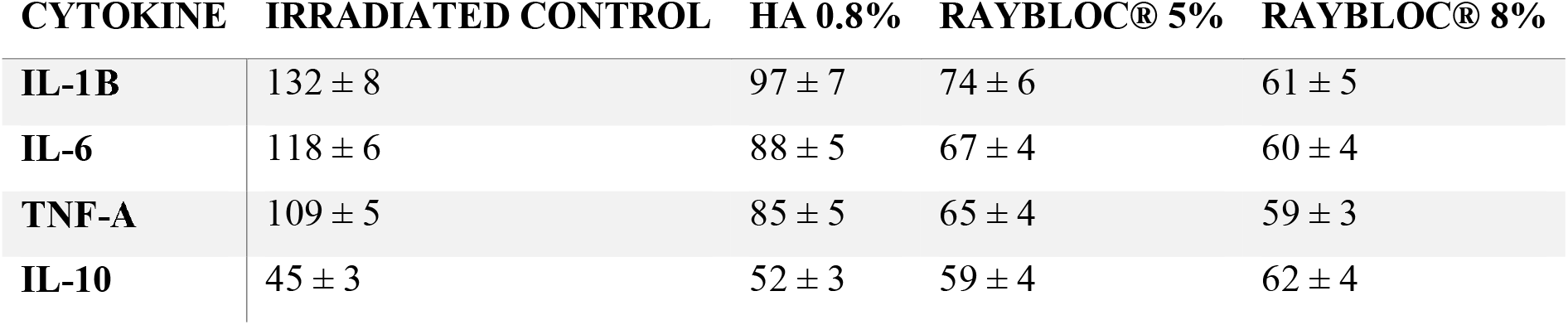
Cytokine levels (pg/mg protein).

| **CYTOKINE** | **IRRADIATED CONTROL** | **HA 0.8%** | **RAYBLOC® 5%** | **RAYBLOC® 8%** |
| --- | --- | --- | --- | --- |
| **IL-1B** | 132 ± 8 | 97 ± 7 | 74 ± 6 | 61 ± 5 |
| **IL-6** | 118 ± 6 | 88 ± 5 | 67 ± 4 | 60 ± 4 |
| **TNF-A** | 109 ± 5 | 85 ± 5 | 65 ± 4 | 59 ± 3 |
| **IL-10** | 45 ± 3 | 52 ± 3 | 59 ± 4 | 62 ± 4 |

## 4. Discussion

This preclinical study shows that Raybloc®, a marine-bioactive silica-microsponge formulation, is effective at protecting skin from damage caused by blue light (HEV) and IR-A. Raybloc® at 8% consistently outperformed hyaluronic acid (HA) comparators across barrier-function, molecular, and histological endpoints, supporting its role as an advanced topical defense system. Raybloc® significantly reduced transepidermal water loss (TEWL) and increased skin hydration, indicating that the barrier was successfully restored. Glycerin and sorbitol are two examples of humectants that work with the sustained-release microsponge matrix to keep actives on the surface longer and hold more water. Microsponge-based systems are well-known for improving the retention of topical agents on the skin, increasing stability, and slowing their diffusion (Shrestha & Banga,2025). Raybloc® bioactive matrix offers greater structural benefits than HA, which primarily works as a moisturizer.

The formulation exhibited significant antioxidant properties, as evidenced by reduced ROS accumulation, MMP-1 inhibition, and activation of the Nrf2–HO-1 cytoprotective pathway. Marine algae extracts, particularly from Chlorella and Laminaria, contain phenolic compounds, carotenoids, peptides, and mycosporine-like amino acids (MAAs) that are known to provide UV/HEV photoprotection through ROS scavenging and Nrf2 pathway activation (Cichoński & Chrzanowski, 2022). The 2.1-fold increase in collagen I and the decrease in MMP-1 are in line with earlier studies showing that marine bioactives can protect the extracellular matrix and slow photoaging (Zhang et al., 2024). Raybloc® also worked well to reduce inflammation by lowering levels of IL-1β, IL-6, and TNF-α and raising levels of IL-10. These results are consistent with what we already know about how marine polysaccharides and peptides can inhibit NF-κB and MAPK inflammatory pathways in skin models exposed to radiation (Chen et al., 2025; Xu & Zhao, 2022). Histological decreases in inflammatory infiltration strengthen Raybloc® effectiveness in reestablishing cutaneous homeostasis following HEV + IR-A exposure.

Raybloc®-treated skin showed significant improvement in collagen preservation and tissue integrity, with about 85 ± 4% of collagen remaining and very little epidermal hyperplasia. Controlled-release microsponge delivery likely ensured that protective antioxidants were always available in the skin, helping protect fibroblasts and the extracellular matrix from damage. Microencapsulated or microsponge-based photoprotection systems have been reported to offer similar benefits (Jadhav et al., 2013; Rahman et al., 2022). Raybloc® provides a more comprehensive approach to UV protection than HA, which is great at keeping skin hydrated but less effective at fighting inflammation and free radicals. Research indicates that multi-component marine-based formulations and systems containing supplementary antioxidants are superior to hyaluronic acid alone in safeguarding against UV and HEV-induced damage (Alves, Sousa, Kijjoa, & Pinto, 2020; Liu, 2022; Sharma, Singh, Srivastava, & Dan, 2025). Raybloc® may be a better choice than regular HA-based products because of the way humectants, marine antioxidants, and microsponge-controlled delivery work together.

Although the preclinical findings are promising, further investigation is required to ascertain their efficacy in humans. Future research should include dose–response evaluations, more detailed mechanistic studies—especially clinical trials with human subjects. The findings indicate that Raybloc® is a promising multifunctional topical formulation capable of safeguarding the skin’s barrier, reducing oxidative and inflammatory stress, and preserving the skin’s structure from damage induced by contemporary light.

## 5. Conclusions

Raybloc®, a topical formulation of marine-bioactive silica-microsponge, showed significant photoprotective effectiveness against HEV blue light and IR-A radiation in a nude mouse model. The formulation significantly improved hydration, collagen preservation, and overall skin structural integrity while effectively reducing transepidermal water loss, oxidative stress, and the expression of inflammatory cytokines. These results show that combining marine antioxidants, humectants, and controlled-release microsponge technology on a single platform can improve outcomes. Raybloc® consistently outperformed hyaluronic acid controls across biophysical, biochemical, and histological parameters, highlighting its multifunctional mechanism of action that transcends traditional moisturization. Raybloc® works through multiple pathways that contribute to skin aging from digital and solar sources by restoring the skin’s barrier, maintaining redox homeostasis, modulating inflammation, and protecting the extracellular matrix. Overall, the results support Raybloc® potential as a new type of skin care product that can help address emerging skin problems in modern light environments.

## Acknowledgments

This work was funded by Singapore Ecommerce Centre Pte Ltd (Singapore), owner of the Nano Skinz brand. Grammarly (pro version) was used to refine the academic language. No AI-generated text was used directly in the final manuscript.

## Competing financial interest

The authors declare the following competing financial interest(s): SY and KN are employees of Singapore Ecommerce Centre Pte Ltd. MO is an employee of PengyouX Pvt Ltd.

